# Blue beats green: Agonistic interactions between Atlantic blue crabs and European green crabs in the Gulf of Maine

**DOI:** 10.1101/2025.09.25.678676

**Authors:** Samantha A. Smith, Reuben P. Siegel, Laura C. Crane, Kayla B. Cheney, Benjamin C. Gutzler, Jaret S. Reblin, Jason S. Goldstein, Christopher D. Wells

## Abstract

As ocean temperatures rise, range expansions and biological invasions are increasingly reshaping marine ecosystems. Warming waters are promoting the northward expansion of the Atlantic blue crab (*Callinectes sapidus*) into the Gulf of Maine (GoM) where its arrival has resulted in habitat overlap and novel interactions with the invasive European green crab (*Carcinus maenas*), a pervasive resident of coastal GoM for over a century. Through a series of laboratory trials, we assessed the interactions of paired blue and green crabs sourced from the GoM, specifically investigating the effects of crab size, aggression, and availability of alternative prey on these interactions. Blue crabs were found to be effective predators of green crabs, killing them in half of trials. Blue crabs were more likely to kill smaller green crabs (< 65 mm carapace width), and small to mid-size blue crabs (< 170 mm carapace width) were more likely than large blue crabs to consume green crabs. There was no green crab predation on blue crabs, though larger green crabs (> 65 mm carapace width) displayed more aggressive behavior towards blue crabs. Presence of an alternative prey item had no effect on blue crab predation on green crabs. Blue crab predation on green crabs could shift future ecosystem dynamics, altering community ecology in the GoM. This study highlights how climate-driven range expansions can mediate interactions among introduced species, with implications for ecosystem structure and species management.

## 1. INTRODUCTION

The introduction of exotic marine species can alter the composition and function of entire ecosystems (Molnar et al., 2008). Novel species can diminish native or existing populations, decrease habitat biodiversity, alter ecosystems, and threaten human health (Bax et al., 2003; Molnar et al., 2008). While anthropogenically driven translocations are historically the leading mechanism of marine species introductions (Molnar et al., 2008), more recent climate-induced changes in temperature (Cheng et al., 2022) and phenology (Edwards and Richardson, 2004) are facilitating the range expansion of species poleward (Lucey and Nye, 2010; Pinsky et al., 2020). Whether through introduction or range-expansion, the establishment of a new species can cause dramatic shifts in ecosystem dynamics and habitat overlap with other species, resulting in novel species interactions (Bax et al., 2003; Gallardo et al., 2016). For example, the invasion of the Asian shore crab (*Hemigrapsus sanguineus*) into the east and west coasts of the United States in the late 1900s spurred competitive interaction and habitat exclusion with the previously established European green crab (*Carcinus maenas*) (Jensen et al., 2002). Novel interactions such as this may multiply as the climate warms and species shift their ranges to accommodate newly habitable territories (Lucey and Nye, 2010).

As sea surface temperature warms rapidly (Pershing, 2018), the western Gulf of Maine (GoM) is experiencing an influx of range-expanding species, altering ecosystem dynamics and species composition (Pershing et al., 2015, 2021; Staudinger et al., 2019; Mills et al., 2024). The GoM is a unique marine ecosystem due to its distinctive positioning between the northern Labrador current and southern Gulf Stream whose circulation patterns influence water temperature and oceanographic conditions and cause broad seasonal variability in the southern GoM (Goode et al., 2019; Lotze et al, 2022; Townsend et al., 2023). These conditions make it a hotspot for climate change impacts and create opportunities to model potential changes in marine ecosystems globally (Lotze et al., 2022). The GoM is historically prone to invasions by non-native species, impacting native populations through both direct (e.g., predation, competition) (Carlton, 2003) and indirect interactions (e.g., habitat exclusion, disease transmission, disruption of ecological interactions) (Crowl et al., 2008; Wagstaff, 2015). Rapid warming exacerbates this vulnerability by accommodating range-expanding species which are now able to establish persistent populations in regions they have not previously inhabited (Lucey and Nye, 2010; Pinsky and Fogarty, 2012; Lawlor et al., 2024). Warming not only catalyzes range expansions but also creates conditions that are conducive for the establishment and persistence of invasive species (Mainka and Howard, 2010).

The invasive European green crab (*C. maenas*, herein green crab) has been a persistent resident in the GoM since introduction via ballast water of ships in the 1800s, and the explosion of its population since the 1940s has had detrimental effects in the GoM (Scattergood, 1952; reviewed in Young & Elliott, 2019). Green crabs are considered highly invasive due to their rapid reproduction rate, ability to tolerate a wide range of environmental conditions, and potential to cause significant and widespread ecological damage to native species and key habitats (Klassen and Locke, 2007; Young and Elliot, 2019; Ens et al., 2022; Wells et al., 2023). In the GoM, green crabs have been shown to have negative ecological and economic impacts on native species and fisheries (Fairchild and Howell, 2000; Griffen and Riley, 2015; Rayner and McGaw, 2019; Quijón, 2024). Green crabs have also been directly implicated in habitat loss through the destruction of eelgrass beds (*Zostera marina*), a vital habitat for native juvenile fishes and shellfish species (Malyshev and Quijón, 2011; Neckles, 2015).

One range-expanding species currently establishing a population in the GoM due to increasingly favorable thermal conditions is the Atlantic blue crab (*Callinectes sapidus*, herein blue crab) (Johnson, 2015; Stasse et al., 2023; Crane et al., 2024). The blue crab is a commercially and ecologically important species that has a historical range from Argentina to the south coast of Cape Cod, Massachusetts (Scattergood, 1960; Johnson, 2015). Along the mid-Atlantic and Gulf Coasts of the United States, the blue crab supports a well-established and economically valuable fishery (Stagg and Whilden, 1997; Miller et al., 2005) and is considered a major predator in its environment (Peterson, 1979; Clavero et al., 2022; Johnson, 2022). Blue crabs are omnivorous generalist predators that typically prey on plant and algal material, fishes, molluscs, and arthropods, including a variety of shellfish and crustaceans (Hines, 2007). Since the 1860s, ephemeral observations of blue crabs have been reported in the GoM, historically restricted to warmer summers as blue crabs were unable to survive winter temperatures (Piers, 1920; Scattergood, 1960). Increases in blue crab sightings in the GoM and as far north as Nova Scotia have typically occurred during abnormally warm years (Piers, 1920; Scattergood, 1960). More recently, in concert with ocean warming (Pershing et al., 2015, 2021), surveys have found a drastic increase in the number of blue crabs in the GoM, with over 350 public blue crab sightings reported in the past five years (GBIF Secretariat, 2023; Marissa McMahan, pers. comm., 2025). Most life stages have now been observed in the GoM, including megalopae, juveniles, adults, mating pairs, gravid females, and overwintering individuals, suggesting persistence of the current population (Stasse et al., 2023; Crane et al., 2024). Though the extent of blue crab population establishment in the GoM remains unclear, the range expansion and possible persistence of this species could dramatically alter local ecosystem dynamics and species composition.

As blue crabs expand their range into the Gulf of Maine, their interactions with green crabs are likely becoming more frequent. Recent observations include blue and green crabs together in the same salt marsh pools and crab traps (Crane, unpub. data, 2025). In baited traps where green crabs are typically numerous, it has been observed that the presence of a blue crab is frequently accompanied by fewer green crabs in the trap (Crane, unpub. data, 2025). The nature of these interactions between blue and green crabs in and around traps in Gulf of Maine is largely unknown, but trap data suggests predation or competition may be a factor between these two species.

Previous studies have investigated agonistic interactions between blue and green crabs and compared their feeding mechanisms and behaviors (de Rivera et al., 2005; MacDonald et al., 2007; Rogers et al., 2018; Davenport et al., 2023). As effective predators of green crabs, blue crabs are thought to limit the abundance and spread of green crabs southward along the Mid-Atlantic coast where blue crabs are native and abundant (de Rivera et al., 2005). However, blue crabs have been found to lose their competitive advantage against green crabs in food contests and exhibit less intraguild predation at temperatures < 16 °C (MacDonald et al., 2007; Rogers et al., 2018). Additionally, juvenile blue crabs were found to be inferior competitors to adult green crabs (MacDonald et al., 2007). These prior studies have found varying levels of blue crab predation on green crabs using specimens collected south of the GoM. There are currently no published studies on the agonistic interactions of adult blue crabs and green crabs in the GoM, though shared habitat and prey preference create the potential for competition and predation. Our study aimed to determine if adult blue crabs in the GoM will prey on adult green crabs under GoM summer water conditions which typically span the 16 °C threshold, unlike their historically inhabited range which stays warmer (Scattergood, 1960). Using a series of laboratory experiments, we quantified blue crab predation rates on green crabs and assessed the effects of relative crab size, aggressiveness, and availability of an alternate prey on their interactions and predation rates. Two trial series were designed and implemented independently of one another at two sites in the GoM, creating the unique and valuable opportunity to compare these complementary studies to more comprehensively investigate blue crab and green crab interactions under varying conditions and at different geographic locations within the blue crab’s expanded range.

## 2. MATERIALS AND METHODS

### 2.1. Study sites and specimen collection

Laboratory trials investigating blue and green crab interactions were conducted at two research stations in the GoM: the Wells National Estuarine Research Reserve (WNERR), Wells, ME and the Schiller Coastal Studies Center (SCSC) of Bowdoin College, Harpswell, ME. Trials at the two locations were developed and conducted independently of one another but have parallel and complementary methodology, providing a unique opportunity to compare results across two study sites in the GoM and expand the scope of the investigation.

In both trial series, blue crabs and green crabs were collected with baited blue crab traps (60 × 50 × 50 cm; Ketcham Supply Co., New Bedford, MA); additional green crabs were collected with baited cylindrical Blanchard-style traps (90 × 40 × 40 cm). For WNERR trials, blue crabs and green crabs were collected from the Little River (43.334 °N, 70.545 °W) and Webhannet River (43.320 °N, 70.564 °W) estuaries, Wells, ME (Fig. 1) in May-August 2024. In SCSC trials, blue crabs were collected in the New Meadows estuary (43.925 °N, 69.864 °W), Bath, ME (Fig. 1) in September-November 2024; green crabs were trapped in Harpswell Sound (43.800°N, 69.953°W), Harpswell, ME (Fig. 1), and the same New Meadows location as the blue crabs.

**Fig. 1.**
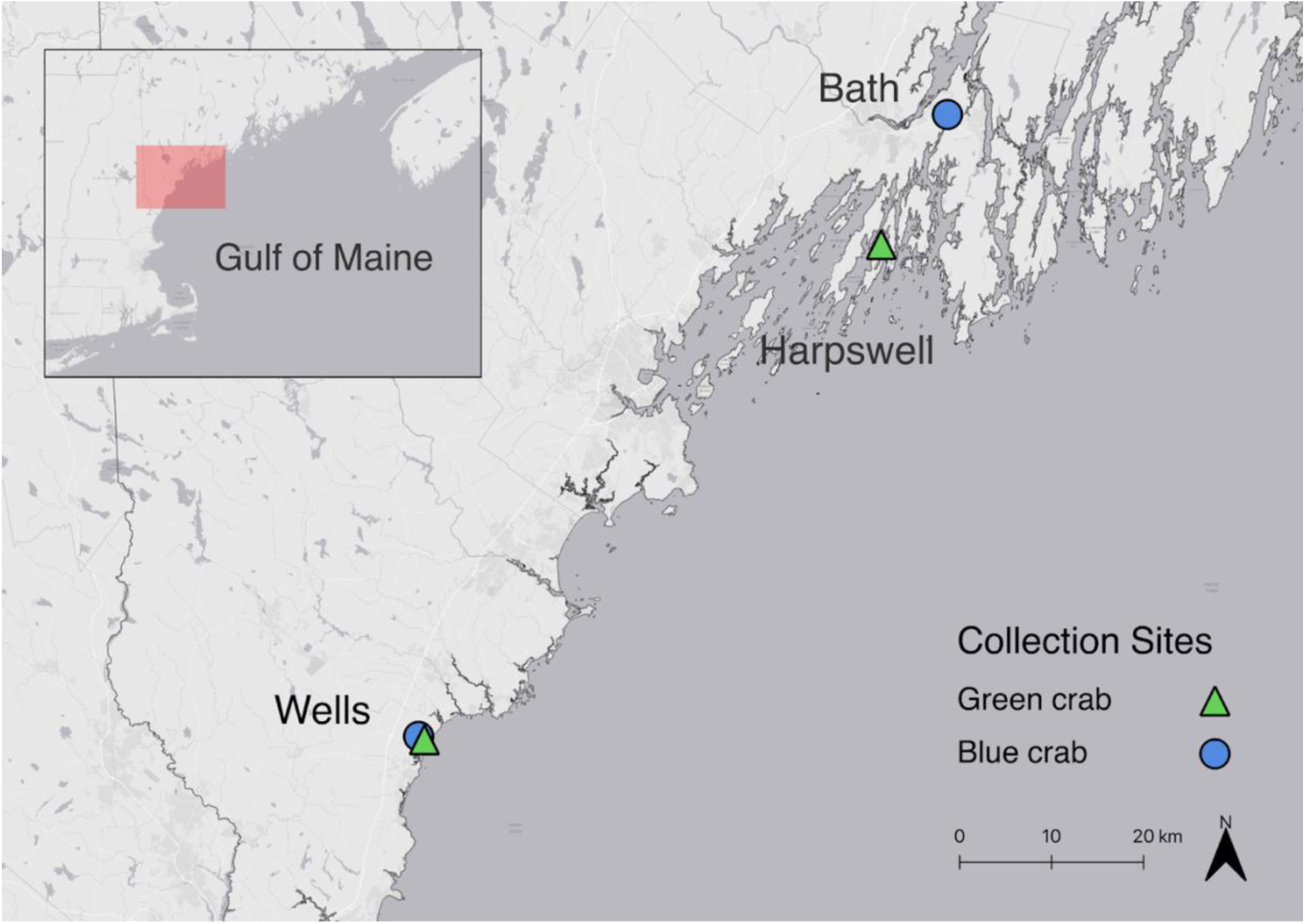
Map of green and blue crab collection sites in the Gulf of Maine.

Crabs at both locations were maintained in aerated aquaria at ambient temperatures (mean = 17.2 °C ± 2.2 SD) and salinities (mean = 31.5 ppt ± 0.5 SD); blue crabs were held individually while green crabs were held communally. Seawater was sourced from the Webhannet River estuary (WNERR) and Harpswell Sound (SCSC). All crabs were weighed (wet weight) and measured (carapace width including lateral spines, herein CW). In this study, we report analyses in relation to CW as the two metrics of size were significantly related in a log-log regression (Fig. S1, n = 45, F_1,43_ ≥ 544, *p* < 0.01). Crabs used in trials had two intact chelipeds and were missing no more than one walking leg. All crabs were fed blue mussels (*Mytilus edulis*) two days prior to a trial and then subsequently starved until the start of the trial. All green crabs were frozen after each trial. At WNERR, blue crabs were only used once and were subsequently tagged and returned to their collection site. At SCSC, due to limited availability of specimens, blue crabs were reused for up to three trials with at least four days in between, during which they were fed mussels daily until the 48 hours prior to trial start. No substratum (i.e., sand) was used as it was determined in pilot trials that sand caused low visibility and did not significantly impact crab interactions. Only male blue crabs were used and both male and female green crabs were used in trials (see Discussion).

### 2.2. Tank trials

To examine interspecific interactions between crabs in a shared habitat, one blue crab and one green crab were paired in 45 trials carried out at WNERR (19 trials) and SCSC (26 trials) using a range of crab size proportions.

At WNERR, trials were conducted in two acrylic recirculating seawater tanks (83 × 83 × 46 cm) with opaque sides to reduce external visual disturbance (Fig. S2). Tanks were filled with 69 L of seawater (10 cm depth). Two bricks were placed in the center of each tank as shelter such that crab sightlines were obstructed but visibility from above was not, with a ceramic tile placed between them. Tank water was filtered using a Penn Plax Cascade Canister (Penn Plax Inc., Hauppauge, NY). Trials were recorded from above with a time lapse camera (Brinno TLC200 Pro, 720P HD, or TLC300, 1080P HD; Brinno Inc., Taipei City, Taiwan) that photographed every 5 seconds. All overhead lighting except a red light (which allowed for night photography) was turned off and natural light was allowed in through two small windows.

At SCSC, trials were conducted in two fiberglass flow-through seawater tanks (57 × 57 × 50 cm) that had three opaque sides and one transparent side through which video was recorded (Fig. S2). The transparent side was covered with an opaque tarp, which enclosed the camera behind the panel, to reduce external visual disturbance and the top was covered with a light diffusing grate. Tanks were filled with approximately 140 L (50 cm of depth) of seawater filtered through 150 micrometer mesh and aerated with an air stone. An opaque PVC sheet rounding the perimeter of each tank blocked access to three of four corners to leave one corner resource available, as preliminary observations showed that both blue and green crabs preferred the corner space. Water temperature was maintained at ∼15 °C using four 800 W titanium heaters. Each trial was recorded for the first four hours with a camera (GoPro Hero 8 Black Edition or Hero 11 Black Edition, wide angle, 30 FPS, 1080P HD) from a side angle (∼30° from horizontal) during which the tank was illuminated with four overhead T5 lights (Agrobrite, 6500 K, 54 W bulbs) which provided lighting for videography and simulated daytime light levels. For the remainder of the trial, overhead fluorescent lighting cycled with the laboratory’s day and night schedule (approximately 9 hours lights-on:15 hours lights-off).

Each trial began with an acclimation period (1 hour at WNERR, 24 hours at SCSC) during which crabs were separated from one another and allowed to acclimate to trial water conditions and ambient chemical cues, including cues from paired trial crabs. Crabs were then allowed to explore their environment freely and interact with each other for the 48-hour trial duration. Trial tanks were cleaned of debris and refreshed with new seawater between trials. At WNERR, green crab to blue crab size ratios ranged from 30-80%; green crabs ranged from 45-71 mm CW (mean = 62 mm ± 9 SD) and blue crabs ranged from 88-155 mm CW (mean = 131 mm ± 17 SD). At SCSC, trials included size proportions of 26-55% green crab to blue crab CW; green crabs were similar in size to those from WNERR (44-81 mm CW, mean = 64 mm ± 11 SD), though blue crabs were larger (135-205 mm CW, mean = 178 mm ± 19 SD).

To test the effect of an alternate food resource on crab interactions, half of the trials (9 of 19) conducted at WNERR included the availability of blue mussels (*Mytilus edulis*) as a prey item. In these trials, ∼400 g of whole, live mussels, each measuring < 3 cm long, were attached with rubber bands to the ceramic tile in the center of the tank one day prior to trial start to allow attachment via byssal threads. Between trials, mussels were removed and tiles were cleaned.

### 2.3. Video and statistical analyses

Video footage of the first four hours of each trial was used to assess crab behavior and interactions. Five behaviors were quantified – intercrab aggression, green crab fatality, green crab consumption, mussel consumption, and corner usage (see Table 1 for definitions). Behavior categories were derived from preliminary observations of blue and green crabs in pilot trials. Green crab consumption, mussel consumption, and corner usage were analyzed as durations of time, whereas intercrab aggression was analyzed as a rate (number of occurrences per hour) and green crab fatality was recorded as a single instance. Aggressive behaviors were counted with respect to the aggressor. Trials were ended following fatal attacks. In the subsequent 44 hours of each trial, crab fatality was noted at 24 and 48 hours, except for the first four trials at SCSC which were terminated after 4 hours.

**Table 1.** Descriptions of behaviors of interest of blue crabs and green crabs, as well as the site(s) at which each behavior was analyzed.

| Category Name | Description | Site Tested |
| --- | --- | --- |
| Intercrab aggression | Crab initiates aggressive behavior (claws raised towards and/or agonistic physical contact with other crab) | Both |
| Green crab fatality | Blue crab fatally attacks green crab | Both |
| Green crab consumption | Blue crab eats green crab tissue (claw movement from green crab to mouth) | Both |
| Mussel consumption | Crab eats mussels (claw movement from mussels to mouth) | WNERR |
| Corner usage | Crab has over half its body in the corner region (Fig. S2) | SCSC |

Data were analyzed in R version 4.5.1 (R Core Team, 2025). Statistical significance was set at α ≤ 0.05. Each trial was considered an independent unit of observation. Blue crabs were used in multiple trials (≤ 3 trials each) at SCSC, although their identity was not tracked, and so blue crab identity was not used as a factor in models. Aggressions per hour corrected for trials where green crabs died before the four-hour recording measurement period had concluded.

We used a binomial regression to assess how CW and aggressiveness of each crab affected green crab survival. CW and aggressions per hour for both species, their two-way interactions, and location were included as fixed effects. Aggressive interactions were modeled as a function of blue and green crab size using a negative binomial regression (R package MASS 7.3-65) (Venables and Ripley, 2002). CW of both species and collection location were included as fixed effects, with an offset term to account for variation in observation time. Trial durations varied as some trials were terminated early when green crabs were killed, precluding further aggression. By estimating aggression rates per hour, both unequal trial lengths and overdispersion were accommodated in the count data. To ensure model convergence, up to 1000 model fitting iterations were allowed.

Competition was assessed in two ways: competition for food and for a corner refuge (i.e., shelter). To assess food-driven competition, we used a binomial generalized linear model to evaluate how the presence of mussels, green and blue crab size (CW), and their interaction influenced green crab survival; only data collected at WNERR were included in this analysis. To assess competition for shelter, the duration of corner use was modeled using a negative binomial generalized linear mixed model (R package glmmTMB 1.1.11) (Brooks et al., 2017), with crab species, crab size (CW), and their interaction as fixed effects; trial number was included as a random effect to account for non-independence within trials; only data collected at SCSC were included in this analysis. Model fitting was permitted up to 1000 iterations to ensure convergence.

## 3. RESULTS

During field collections, blue and green crabs were captured together in the same traps, indicating natural co-occurrence at our study sites.

### 3.1. Predation

Blue crabs killed green crabs in 49% of trials (20 of 41 trials) and green crabs never killed blue crabs. Several factors influenced green crab survival. The effect of green crab size on survival varied with blue crab size (Fig. 2A, binomial regression, χ²_1,31_ = 5.6, *p* = 0.02). Blue crabs were more likely to prey on smaller green crabs, and larger blue crabs were less likely to prey on green crabs regardless of their size. Additionally, green crab CW interacted significantly with green crab aggression (Fig. 2B, binomial regression, χ²_1,31_ = 4.4, *p* = 0.04). Larger green crabs (> 65 mm CW) were more likely to survive than smaller green crabs, but green crabs that exhibited little to no aggression were less likely to survive regardless of their carapace width. The aggressiveness of blue crabs also significantly affected the survival of green crabs with more aggressive blue crabs more likely to kill green crabs (Fig. 2C, χ²_1,31_ = 5.3, *p* = 0.02). Location of collection did not significantly affect green crab survival (χ²_1,31_ = 0.5, *p* = 0.52).

**Fig. 2.**
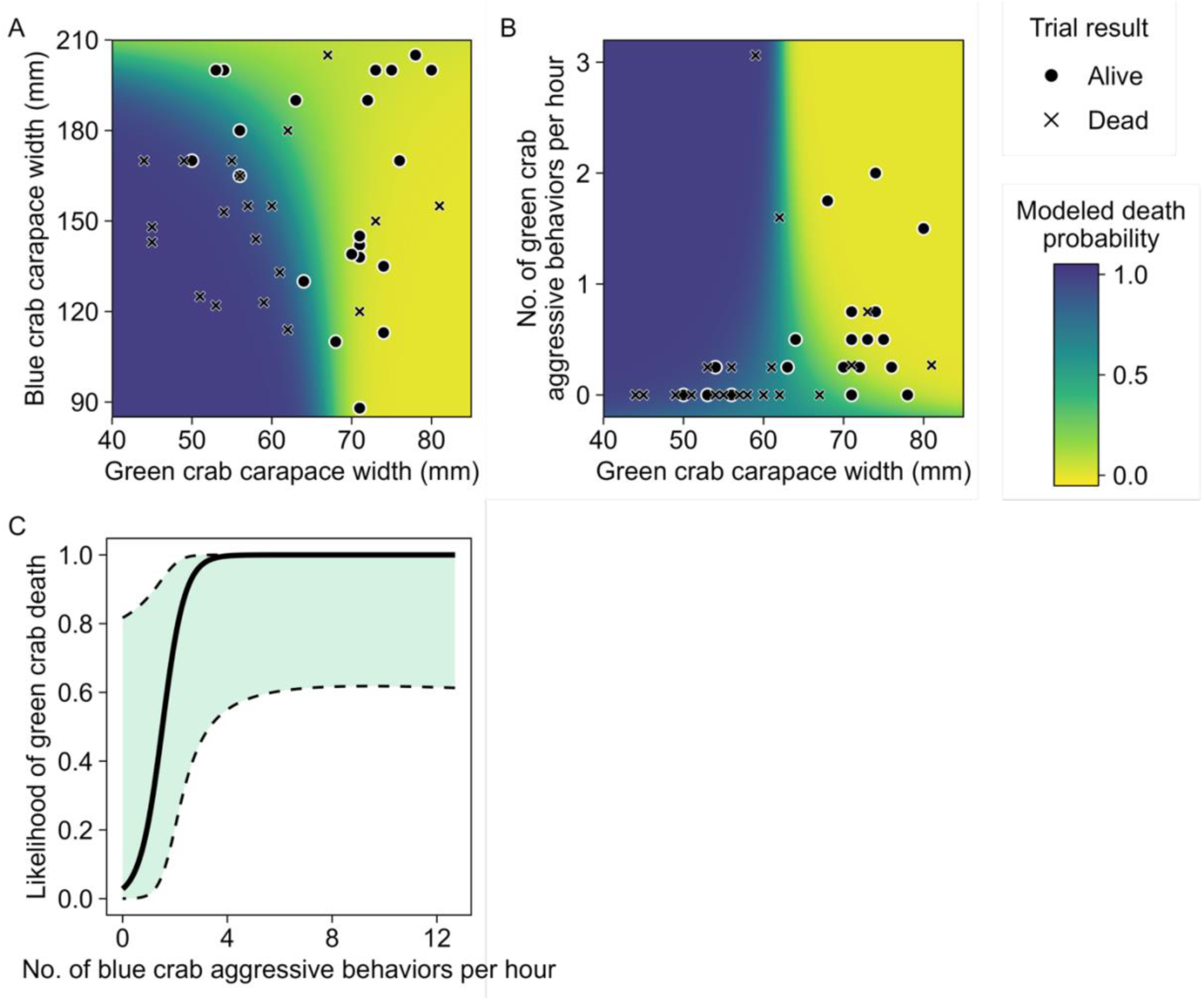
Green crab mortality as a function of (A) green crab and blue crab carapace width (CW), (B) green crab CW and green crab aggression, and (C) blue crab aggression. (A, B) Heatmaps display the modeled probability of green crab mortality with purple indicating high predicted mortality and yellow indicating low predicted mortality. Points indicate observed trial outcomes, with circles representing green crabs surviving for the 48-h trial duration and crosses denoting green crab mortality. (C) The bold line represents the model prediction, while the shaded region bound by dashed lines indicates the 95% confidence interval.

### 3.2. Aggression

Green crab aggression towards blue crabs was significantly affected by green crab CW (Fig. 3, negative binomial regression, χ²_1,37_ = 17.6, *p* < 0.01), but not by blue crab size nor their collection location (negative binomial regression, χ²_1,31_ ≤ 2.61, *p* ≥ 0.10). Green crabs were more aggressive when they were larger; a 1 mm increase in green crab CW corresponded with an approximate 9.7% increase in green crab aggression rate. In the first four hours of a trial, large green crabs (> 65 mm CW) initiated an average of 0.65 aggressions per hour (± 0.68 SD) while small green crabs initiated an average of 0.27 aggressions per hour (± 0.62 SD). Blue crabs initiated an average of 1.32 aggressions per hour (± 1.14 SD). Blue crab aggression was independent of blue crab CW or green crab CW (negative binomial regression, χ² ≤ 2.7, *p* ≥ 0.10).

**Fig. 3.**
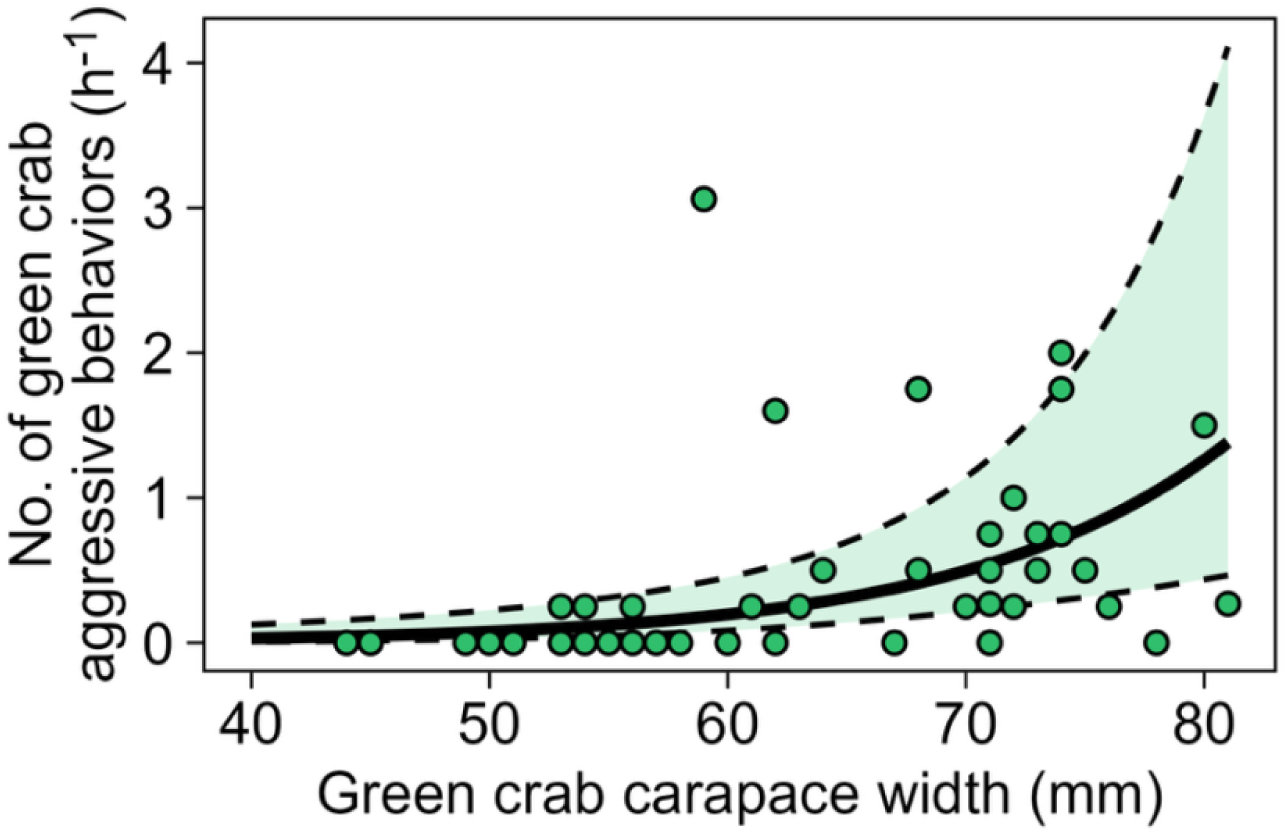
Predicted probability of green crab aggression towards blue crabs as a function of green crab carapace width. The bold line represents the model prediction, while the shaded region bound by dashed lines indicates the 95% confidence interval. Points indicate observed aggression rates.

Many interactions between blue and green crabs were nonaggressive. Crabs would occasionally sit in proximity or on top of each other, with no sign of agonism. Blue crabs spent hours of each trial walking or swimming back and forth along a wall of the tank. Blue crabs often cleaned debris off their chelipeds using their front walking legs. Green crabs moved more sporadically around the tank than blue crabs. When blue crabs attacked green crabs, they often raised themselves onto all walking legs with a meral spread (i.e., claws outstretched laterally). Many blue crab predation events occurred as the result of a green crab approaching a blue crab.

### 3.3. Competition

The presence of mussels did not have a significant impact on green crab fatality (binomial regression, χ²_1,14_ = 3.6, *p* = 0.06). As in the whole dataset, green crab death in trials with mussels present was significantly dependent upon green crab CW, with larger green crabs being less likely to die (binomial regression, χ²_1,14_ = 17.5, *p* < 0.01). Blue crabs consumed green crabs in 67% of trials with mussels and 50% of trials without mussels. In trials with mussels, blue crabs consumed mussels before killing a green crab in 45% of trials, after which blue crabs killed green crabs 25% of the time. In trials where blue crabs killed green crabs, the blue crab consumed the green crab before consuming mussels 83% of the time (5 of 6 trials). Green crabs consumed mussels before blue crabs consumed mussels 78% of the time (7 of 9 trials), excluding one trial in which the green crab was killed within seconds of trial start and thus had no chance to eat mussels. In addition, green crabs began consuming mussels significantly earlier in the trial than blue crabs (mean elapsed time until initial mussel consumption = 1 h 32 min for green crabs, 13 h 11 min for blue crabs).

The amount of time crabs spent in the corner refuge was not related to crab species nor CW (negative binomial generalized linear mixed model, χ²_1,22_ ≤ 0.13, *p* ≥ 0.72). On average, blue crabs spent 22.8% of their time (55 min of 4 h) in the corner refuge whereas green crabs spent 29.4% of their time (1 h 11 min of 4 h) in the corner.

## DISCUSSION

Our findings that blue crabs are effective predators of green crabs augment the conclusions of previous studies (de Rivera et al., 2005; Rogers et al., 2018) by examining interactions in typical GoM summer conditions and investigating the role of aggression, size, and food availability in blue crab predation on green crabs. Like de Rivera et al. (2005), we found that predation likelihood increased when green crabs were smaller (< 67 mm CW). Additionally, we found that large blue crabs (> 170 mm CW) exhibited a lower likelihood of consuming green crabs. Likewise, blue crabs were more likely to prey upon small, less aggressive green crabs than large, more aggressive green crabs. Notably, we did not observe any green crab predation on blue crabs. However, previous studies have found that adult green crabs (55-80 mm CW) are able to kill juvenile blue crabs (< 80 mm) (MacDonald et al., 2007). This vulnerability may impact blue crab persistence in the GoM if green crab predation causes mortality of juvenile blue crabs before they are able to reach a size refuge (Rogers et al., 2018).

Temperature plays a crucial role in the behavior and survival of blue crabs in the GoM and has the potential to moderate interactions between blue and green crabs (de Rivera et al., 2005; Rogers et al., 2018). Prior research has found that blue crab predation on green crabs increases at higher temperatures and significantly drops off at temperatures < 16 °C (Rogers et al., 2018; Connolly-Randazzo, 2022). The temperature range of our trials (15-20 °C) spans this threshold and approximates typical GoM summer sea surface temperatures (Birkel, 2025) simulating blue and green crab interactions under GoM conditions. Ongoing warming climate trends in the GoM may significantly affect the dynamics between these two species and determine which crab species holds a competitive advantage, potentially determining future patterns of dominance and displacement in the GoM.

A prediction of optimal foraging theory is that predators prefer smaller and less dangerous prey (Smallegange et al., 2008). Our study showed that blue crabs preferred smaller, less aggressive green crabs, but did not demonstrate a preference for mussels when available. Unlike mussels, green crabs can inflict defensive damage (e.g., appendage removal, carapace breakage) which makes them a high-risk prey item (Sneddon et al., 2000). Blue crabs, among other organisms, have been shown to alter their foraging behavior in the presence of aggressive conspecifics, minimizing conflict and avoiding risk of injury (Abrahams and Dill, 1989; reviewed in Juanes and Smith, 1995; Clark et al., 1999a), which we observed in interspecific interactions. Large, aggressive green crabs may pose an unacceptable risk to blue crabs, potentially explaining the lower predation rates on these individuals (Mukherjee and Heithaus, 2013). Our findings showed that blue crab aggression also played a role in green crab mortality, with more aggressive blue crabs more likely to prey on green crabs. Previous studies have found that blue crab aggression may play an important role in conspecific interactions with similar-sized individuals but hold less importance in interactions with crabs of a greater size difference (Reichmuth et al., 2011). This suggests that aggressiveness of both blue and green crabs is more important when the two are closer in size. It is likely that the combination of both size and aggressiveness of the two individuals is responsible for the outcome of the interaction, as demonstrated in our study. Blue crabs have historically provided tests of optimal foraging theories (Clark et al., 2000) and our study provides further reinforcement of their value in testing these models.

In interspecific food competition trials, green crabs were quicker to find mussels than blue crabs, corroborating previous studies (de Rivera et al., 2005; MacDonald et al., 2007; Rogers et al., 2018). This finding is supported by differences in green and blue crab morphology, as the shape of green crab chelae make them ideal for crushing durable prey, while the shape of blue crab chelae make them more suited for speedy attacks but have lower crushing force (Schenk and Wainwright, 2001). Conversely, the diet of blue crabs in Chesapeake Bay primarily consists of bivalve molluscs (60%), while arthropods, including crabs, comprise a smaller percentage (Hines, 2007). In our trials, blue crabs showed no preference for mussels over green crabs and readily preyed on green crabs even when given an alternative option, suggesting that blue crabs will consume green crabs even in an environment with other prey options available, such as the GoM.

While blue crabs may offer some natural predation pressure, human-mediated efforts to control green crab populations have faced significant obstacles. Logistical barriers such as low economic value, minimal market infrastructure, and preference for recently molted soft-shell crabs have so far limited the establishment of green crab fisheries in their invaded ranges (Walter, 2021). While trapping has been proposed as a mechanism for the control of some green crab populations, more efforts have been directed at mitigation (reviewed by Ens et al., 2022). However, with blue crabs expanding northward and establishing a persistent population, it is possible that blue crabs may provide a biological control to the green crab population in the GoM, particularly in soft bottom and brackish environments where blue crabs have been observed (de Rivera et al., 2005; Johnson, 2015; Prior et al., 2018; Crane et al., 2024). The control of green crabs may allow for a recovery of impacted eelgrass beds and shellfish populations, which could improve water quality and reduce shoreline erosion (Malyshev and Quijón, 2011; Neckles, 2015). It is important to note that blue crabs are opportunistic feeders (Hines, 2007), and their impacts on other populations in the GoM, including shellfish, are unknown. However, a sustained population of blue crabs in the GoM could have ecological and economic benefits, by offering some degree of biological control of green crabs (Lafferty and Kuris, 1996; de Rivera et al., 2005; Prior et al., 2018) and supplementing the economically valuable American lobster (*Homarus americanus*) industry which faces temperature-mediated declines (Rheuban et al., 2017; Jarrett et al., 2024). As blue crabs become more abundant in the GoM, their generalist diet raises concerns about potential negative interactions with existing shellfish and lobster fisheries, particularly if predation pressure shifts toward commercially valuable species.

There are several gaps that remain in understanding the ecological interactions between these two species. First, all studies, including our own, have examined the interactions between these two species on a one-on-one basis (de Rivera et al., 2005; MacDonald et al., 2007; Rogers et al., 2018). Studies replicating blue crab interactions with multiple green and blue crabs would better approximate natural density dynamics, especially given the high density of green crabs in the GoM (Webber, 2013) and the cannibalistic nature of blue crabs and green crabs (Clark et al., 1999b; Gehrels et al., 2017). In our trapping efforts, we collected blue and green crabs together in the same traps, suggesting that such co-occurrence is possible under natural conditions and underscoring the importance of testing interactions under more realistic multi-individual scenarios. Second, understanding larval dispersal mechanisms and blue crab growth rates in their expanded range in the GoM will be helpful, especially since blue crabs have a 4-5 week larval planktonic phase during which they are highly vulnerable to predation (Johnson and Hester, 1989; Epifanio, 1995). An inability to persist through larval, post-larval, and juvenile life stages could limit their establishment in the GoM, thus minimizing the significance of their adult interactions with green crabs. Third, as blue crabs are a novel species in the GoM, little is known about their diet preferences in the region. Examinations of stomach contents, stable isotope studies, and metagenomic analyses may provide valuable insight into their local diet. For example, global studies have found crabs to be the second or third most abundant food category in blue crab stomachs, only after mollusks and fishes (Rady et al., 2018; Taylor et al., 2022; Ortega-Jiménez et al., 2024). However, a recent study in Great Bay Estuary, New Hampshire, found no evidence of green crab in blue crab stomach content analysis (Meyer-Rust et al., 2024).

Lastly, our study was limited by seasonal availability and the size and sex of trapped blue crabs, with most trapped specimens being adult males (≥ 88 mm CW), reflecting the expected sex ratio in lower salinity environments (Marchessaux et al., 2023) where we collected blue crabs. The use of male blue crabs, which are known to exhibit more intraspecific aggression and have proportionally longer chelipeds than females (Jivoff, 1997), may have resulted in more aggressions and interspecific predation in our trials than if we had used female blue crabs. The reuse of individual blue crabs in multiple trials, due to limited catch, may have introduced habituation effects which could have resulted in higher or lower predation rates. However, blue crabs have been found to display less habituation to given environments than other crab species (Balcı et al., 2014). Overall, expanding studies to include use of juvenile blue crabs, female blue crabs, and a broader seasonal window and temperature range could provide further insight into the nuances of blue crab predation on green crabs.

This study focuses on the effects of green crab size and aggression levels, rather than sex, on the likelihood a given green crab will be preyed on by a blue crab. A mix of mostly male and some female green crabs were used in this study, based on what was available at our study sites. Green crabs exhibit sexual dimorphism in cheliped morphology and maximum size, as males tend to be larger and exhibit more aggressive behavior than females (Kattler et al., 2023). In this study, we show that large, aggressive green crabs are less likely to be preyed on by blue crabs. Therefore, we suspect that blue crab predation on green crabs may be higher on female versus male green crabs. However, this was not a focus of the present study. By using a mix of male and female green crabs, we more accurately reflect the population of green crabs that blue crabs will encounter in the natural environment.

In summary, our study highlights the complex size- and behavior-dependent nature of blue and green crab interactions. More broadly, this work contributes to our understanding of predator–prey dynamics and interspecific competition in systems experiencing biogeographic shifts. As climate change and anthropogenic activities continue to facilitate the movement of species beyond their historic ranges, novel interactions between native, invasive, and range-expanding species are becoming increasingly common. Understanding the outcomes of these interactions is critical for predicting ecological impacts, managing invasive and range-expanding species, and anticipating changes in community structure under ongoing environmental change.

## ACKNOWLEDGEMENTS

Thank you to the staff, volunteers, and students at the Wells National Estuarine Research Reserve, the Bowdoin College Schiller Coastal Studies Center and Biology Department, and the Manomet Conservation Sciences for your help with crab trapping, laboratory assistance, and logistical support. Special appreciation for the efforts of Jessie Batchelder, Ruxin Dai, Hadley Horch, Heidi Franklin, Holly Parker, Rachel Reuling, Gordon Shannon, Layla Silva, and Clint Thompson. Trapping at WNERR was conducted under ME-DMR SL-2024-09-04 and trapping at SCSC was conducted under ME-DMR SL-2024-11-04 EDU. Support for this project was awarded from Maine Sea Grant Award NA24OARX417C0036-T1-01, NOAA OPS Award NA23NOS4200175, Peter J. Grua and Mary G. O’Connell Student Research Fund, and the Bowdoin College Biology Department Student Research Fund.

## LITERATURE CITED

Abrahams, M., Dill, L., 1989. A determination of the energetic equivalence of the risk of predation. Ecology 70, 999–1007.

Balcı, F., Balcı-Ramey, P., Ruamps, P., 2014. Spontaneous alternation and locomotor activity in three species of marine crabs: green crab (*Carcinus maenas*), blue crab (*Callinectes sapidus*), and fiddler crab (*Uca pugnax*). Journal of Comparative Psychology 128, 65–73.

Bax, N., Williamson, A.M., Aguero, M., Gonzalez, E., Geeves, W., 2003. Marine invasive alien species: a threat to global biodiversity. Marine Policy 27, 313–323.

Birkel, S., 2025. Gulf of Maine Daily Sea Surface Temperatures. Maine Climate Office, University of Maine.

Brooks, M., Kristensen, K., van Benthem, K., Magnusson, A., Berg, C., Nielsen, A., Skaug, H., Mächler, M., Bolker, B., 2017. glmmTMB balances speed and flexibility among packages for zero-inflated generalized linear mixed modeling. The R Journal 9, 378–400.

Carlton, J., 2003. Community assembly and historical biogeography in the North Atlantic Ocean: the potential role of human-mediated dispersal vectors. Hydrobiologia 503, 1–8.

Cheng, L., von Schuckmann, K., Abraham, J., Trenberth, K., Mann, M., Zanna, L., England, M., Zika, J., Fasullo, J., Yu, Y., Pan, Y., Zhu, J., Newsom, E., Bronselaer, B., Lin, X., 2022. Past and future ocean warming. Nature Reviews Earth & Environment 3, 776–794.

Clark, M., Wolcott, T., Wolcott, D., Hines, A., 1999a. Intraspecific interference among foraging blue crabs Callinectes sapidus: interactive effects of predator density and prey patch distribution. Marine Ecology Progress Series 178, 69–78.

Clark, M., Wolcott, T., Wolcott, D., Hines, A., 1999b. Foraging and agonistic activity co-occur in free-ranging blue crabs (*Callinectes sapidus*): Observation of animals by ultrasonic telemetry. Journal of Experimental Marine Biology and Ecology 233, 143–160.

Clark, M., Wolcott, T., Wolcott, D., Hines, A., 2000. Foraging behavior of an estuarine predator, the blue crab Callinectes sapidus in a patchy environment. Marine Pollution Bulletin 176, 113479.

Clavero, M., Franch, N., Bernardo-Madrid, R., Lopez, V., Abello, P., Queral, J.M., Mancinelli, G., 2022. Severe, rapid and widespread impacts of an Atlantic blue crab invasion. Marine Pollution Bulletin 176, 113479.

Connolly-Randazzo, E., 2022. Temperature and predator effects on green crabs (*Carcinus maenas*) and their distribution in South Slough National Estuarine Research Reserve Environmental Science and Management. Portland State, Portland, Oregon.

Crane, L.C., Burke, E.A., Gutzler, B.C., Goldstein, J., 2024. Evidence of a blue crab (*Callinectes sapidus*) successfully overwintering in a Southern Maine salt marsh. Northeastern Naturalist 31, N11–N16.

Crowl, T.A., Crist, T., Parmenter, R., Belovsky, G., Lugo, A., 2008. The spread of invasive species and infectious disease as drivers of ecosystem change Frontiers in Ecology and Evolution 6, 238–246.

Davenport, J., Jessopp, M., Harman, L., Micaroni, V., McAllen, R., 2023. Feeding, agonistic and cooperative behavioural responses of shallow-water benthic marine scavengers. Natural History 57, 1049–1065.

De Rivera, C., Ruiz, G.M., Hines, A.H., Jivoff, P., 2005. Biotic resistance to invasion: Native predator limits abundance and distribution of an introduced crab. Ecology 86, 3364–3376.

Edwards, M., Richardson, A., 2004. Impact of climate change on marine pelagic phenology and trophic mismatch. Nature 430, 881–884.

Ens, N., Harvey, B., Davies, M., THomson, H., Meyers, K., Yakimishyn, J., Lee, L., McCord, M., Gerwing, T., 2022. The Green Wave: reviewing the environmental impacts of the invasive European green crab (*Carcinus maenas*) and potential management approaches. Environmental Reviews 30, 306–322.

Epifanio, C.E., 1995. Transport of blue crab (*Callinectes sapidus*) larvae in the waters off Mid-Atlantic States. Bulletin of Marine Science 57, 713–725.

Fairchild, E., Howell, W., 2000. Predator-prey size relationship between *Pseudopleuronectes americanus* and *Carcinus maenas*. Journal of Sea Research 44, 81–90.

Gallardo, B., Clavero, M., Sánchez, M., Vilà, M., 2016. Global ecological impacts of invasive species in aquatic ecosystems. Global Change Biology 22, 151–163.

GBIF Secretariat, 2023. *Callinectes sapidus* Rathburn, 1896, in: Taxonomy, G.B. (Ed.).

Gehrels, H., Flynn, P., Cox, R., Quijón, P., 2017. Effects of habitat complexity on cannibalism rates in European green crabs (Carcinus maenas Linnaeus, 1758). Marine Ecology 38, e12448.

Goode, A.G., Brady, D.C., Steneck, R.S., Wahle, R.A., 2019. The brighter side of climate change: how local oceanography amplified a lobster boom in the Gulf of Maine. Global Change Biology 25, 3906–3917.

Griffen, B.D., Riley, M., 2015. Potential impacts of invasive crabs on one life history strategy of native rock crabs in the Gulf of Maine. Biological Invasions 17, 2533–2544.

Hines, A., 2007. Ecology of juvenile and adult blue crabs, in: Kennedy, V., Cronin, L. (Eds.), The Blue Crab: Callinectes sapidus. Maryland Sea Grant College.

Hughes, R., Seed, R., 1995. Behavioural mechanisms of prey selection in crabs. Journal of Experimental Marine Biology and Ecology 193, 225–238.

Jarrett, R., Brady, D.C., Wahle, R.A., Steneck, R., 2024. Shifts in habitat use and demography of American lobsters in coastal Maine (USA) over the past quarter century. Marine Ecology Progress Series 746, 87–98.

Jensen, G.C., McDonald, P.S., Armstrong, D.A., 2002. East meets west: competitive interactions between green crab Carcinus maenas, and native and introduced shore crab Hemigrapsus spp. Marine Ecology Progress Series 225, 251–262.

Jivoff, P., 1997. Sexual competition among male blue crab, *Callinectes sapidus*. Biological Bulletin 193, 368–380.

Johnson, D., 2015. The savory swimmer swims North: A Northern range expansion of the blue crab *Callinectes sapidus*. Journal of Crustacean Biology 35, 105–110.

Johnson, D., 2022. Beautiful swimmers attack at low tide. Ecology 103, e3787.

Johnson, D., Hester, B., 1989. Larval transport and its association with recruitment of blue crabs to Chesapeake Bay. Estuarine, Coastal and Shelf Science 28, 459–472.

Juanes, F., Smith, L.D., 1995. The ecological consequences of limb damage and loss in decapod crustaceans: a review and prospectus. J. Experimental Marine Biology and Ecology 193, 197–223.

Kattler, K., Oishi, E., Lim, E., Watkins, H., Côté, I., 2023. Functional responses of male and female European green crabs suggest potential sex-specific impacts of invasion. PeerJ 11, e15424.

Klassen, G., Locke, A., 2007. A biological synopsis of the European green crab, *Carcinus maenas*. Canadian Manuscript Report of Fisheries and Aquatic Sciences 2818.

Lafferty, K.D., Kuris, A., 1996. Biological control of marine pests. Ecology 77, 1989–2000.

Lawlor, J., Comte, L., Grenouillet, G., Lenoir, J., Baecher, A., Bandara, R., Bertrand, R., Chen, I.-C., Diamond, S., Lancaster, L., 2024. Mechanisms, detection and impacts of species redistributions under climate change. Nature Reviews Earth & Environment 5, 351–368.

Lotze, H.K., Mellon, S., Coyne, J., Betts, M., Burchell, M., Fennel, K., Dusseault, M., Fuller, S., Galbraith, E.D., Suarez, L., de Gelleke, L., Golombek, N., Kelly, B., Kuehn, S., Oliver, E.C.J., MacKinnon, M., Muraoka, W., Predham, W., Rutherford, K., Shackell, N.L., Sherwood, O., Sibert, E., Kienast, M., 2022. Long-term ocean and resource dynamics in a hotspot of climate change. FACETS 7, 1142–1184.

Lucey, S.M., Nye, J., 2010. Shifting species assemblages in the Northeast US Continental Shelf Large Marine Ecosystem. Marine Ecology Progress Series 415, 23–33.

MacDonald, J., Roudez, R., Glover, T., Weis, J., 2007. The invasive green crab and Japanese shore crab: behavioral interactions with a native crab species, the blue crab. Biological Invasions 9, 837–848.

Mainka, S., Howard, G., 2010. Climate change and invasive species: double jeopardy. Integrative Zoology 5, 102–111.

Malyshev, A., Quijón, P.A., 2011. Disruption of essential habitat by a coastal invader: new evidence of the effects of green crabs on eelgrass beds. ICES Journal of Marine Science 68, 1852–1856.

Marchessaux, G., GjoniI, V., Sarà, G., 2023. Environmental drivers of size-based population structure, sexual maturity and fecundity: A study of the invasive blue crab *Callinectes sapidus* (Rathbun, 1896) in the Mediterranean Sea. PLoS ONE 18, e0289611.

Meyer-Rust, K., Strickland, A., Lee, B.-Y., Sevigny, J., Bradt, G., Brown, B., 2024. Diet of the blue crab (*Callinectes sapidus*) during range expansion in Great Bay Estuary, New Hampshire. BMC Genomics 25, 1238.

Miller, T., Martell, S., Bunnel, F., Bonzek, C., Hewitt, D., Hoenig, J.M., Lipcius, R.N., 2005. Stock assessment of the blue crab in Chesapeake Bay 2005: Final report UMCES technical series. Virginia Institute of Marine Science, William & Mary, Virginia, USA.

Mills, K.E., Kemberling, A., Kerr, L.A., Lucey, S.M., McBride, R.S., Nye, J.A., Pershing, A.J., Barajas, M., Lovas, C.S., 2024. Multispecies population-scale emergence of climate change signals in an ocean warming hotspot. ICES Journal of Marine Science 81, 375–389.

Molnar, J., Gamboa, R., Revenga, C., Spalding, M., 2008. Assessing the global threat of invasive species to marine biodiversity. Frontiers in Ecology and the Environment 6, 485–492.

Mukherjee, S., Heithaus, M., 2013. Dangerous prey and daring predators: a review. Biological Reviews 88, 550–563.

Neckles, H.A., 2015. Loss of eelgrass in casco Bay, Maine, linked to green crab disturbance. Northeast Naturalist 22, 478–500.

Ortega-Jiménez, E., Cuesta, J., Laiz, I., González-Ortegón, E., 2024. Diet of the Invasive Atlantic Blue Crab *Callinectes sapidus* Rathbun, 1896 (Decapoda, Portunidae) in the Guadalquivir Estuary (Spain). Estuaries and Coasts, 1075–1085.

Pershing, A.J., Alexander, M.A., Brady, D.C., Brickman, D., Curchitser, E.N., Diamond, A.W., McClenachan, L., Mills, K.E., Nichols, O.C., Pendleton, D.E., Record, N.R., Scott, J.D., Staudinger, M.D., Wang, Y., 2021. Climate impacts on the Gulf of Maine ecosystem: A review of observed and expected changes in 2050 from rising temperatures. Elementa: Science of the Anthropocene 9, 00076.

Pershing, A.J., Alexander, M.A., Hernandez, C.M., Kerr, L.A., Le Bris, A., Mills, K.E., Nye, J.A., Record, N.R., Scannell, H.A., Scott, J.D., Sherwood, G.D., Thomas, A.C., 2015. Slow adaptation in the face of rapid warming leads to collapse of the Gulf of Maine cod fishery. Science 350, 809–812.

Pershing, A.J., Mills, K., Dayton, A., Franklin, B.S., Kennedy, B.T., 2018. Evidence for adaptation from the 2016 marine heatwave in the Northwest Atlantic Ocean. Oceanography 31, 152–161.

Peterson, C.H., 1979. The importance of predation and competition in organizing the intertidal epifaunal communities of Barnegat Inlet, New Jersey. Oecologia 39, 1–24.

Piers, H., 1920. The blue crab (Callinectes sapidus Rathbun): Extension of its range northward to near Halifax, Nova Scotia. Proceedings of the Nova Scotia Institute of Science 15, 83–90.

Pinsky, M., Selden, R.L., Kitchel, Z.J., 2020. Climate-driven shifts in marine species ranges: Scaling from organisms to communities. Annual Review of Marine Science 12, 153–179.

Pinsky, M.L., Fogarty, M.J., 2012. Lagged social-ecological responses to climate and range shifts in fisheries. Climatic Change 115, 883–891.

Prior, K., Adams, D., Klepzig, K., Hulcr, J., 2018. When does invasive species removal lead to ecological recovery? Implications for management success. Biological Invasions 20, 267–283.

Quijón, P.A., 2024. Predator-prey interactions in a coastal setting: Linking crab feeding rates to small scale distribution of clams. Marine Environmental Research 196, 106452.

R Core Team, 2025. R: A language and environment for statistical computing, R Foundation for Statistical Computing, Vienna.

Rady, A., Sallam, W., Abdou, N., El-Sayed, A., 2018. Food and feeding habits of the blue crab, *Callinectes sapidus* (Crustacea: Decapoda: Portunidae) with special reference to the gastric mill structure. Egyptian Journal of Aquatic Biology & Fisheries 22, 417–431.

Rayner, G., McGaw, I.J., 2019. Effects of the invasive green crab (*Carcinus maenas*) on American lobster (*Homarus americanus*): food acquisition and trapping behaviour. Journal of Sea Research 144, 95–104.

Rheuben, J., Kavanaugh, M.T., Doney, S.C., 2017. Implications of future Northwest Atlantic bottom temperatures on the American lobster (*Homarus americanus)* fishery. Journal of Geophysical Research: Oceans 122, 9387–9398.

Rogers, T., Gouhier, T., Kimbro, D., 2018. Temperature dependency of intraguild predation between native and invasive crabs. Ecology 99, 885–895.

Scattergood, L.W., 1952. The distribution of the green crab, *Carcinides maenas* (L.) in the Northwestern Atlantic. 1952, Marine Resources Documents. Maine Department of Marine Resources.

Scattergood, L.W., 1960. Blue crabs (*Callinectes sapidus*) in Maine. Maine Field Naturalist 16, 59–63.

Schenk, S., Wainwright, P., 2001. Dimorphism and the functional basis of claw strength in six brachyuran crabs. Journal of Zoology 225, 105–119.

Smallegange, I., Hidding, B., Eppenga, J., van der Meer, J., 2008. Optimal foraging and risk of claw damage: How flexible are shore crabs in their prey size selectivity? Experimental Marine Biology and Ecology 367, 157–163.

Sneddon, L.U., Huntingford, F., Taylor, A., Orr, J., 2000. Weapon strength and competitive success in the fights of shore crabs (*Carcinus maenas*). Zoology 250, 397–403.

Stagg, C., Whilden, M., 1997. The history of Chesapeake Bay’s blue crab (*Callinectes sapidus)*: fisheries and management. Investigaciones marinas 25, 93–104.

Stasse, A., Meyer, K., Williams, E., Bradt, G., Brown, B., 2023. First documentation of mating blue crabs, *Callinectes sapidus*, in Great Bay Estuary, New Hampshire. Northeastern Naturalist 30, N8–N12.

Staudinger, M.D., Mills, K.E., Stamieszkin, K., Record, N.R., Hudak, C.A., Allyn, A.J., Diamond, A.W., Friedland, K.D., Golet, W., Henderson, M.E., Hernandez, C.M., Huntington, T.G., Ji, R., Johnson, C.L., Johnson, D., Jordaan, A., Kocik, J., Li, Y., Liebman, M., Nichols, O.C., Pendleton, D.E., Richards, R.A., Robben, T., Thomas, A.C., Walsh, H.J., Yakola, K., 2019. It’s about time: A synthesis of changing phenology in the Gulf of Maine ecosystem. Fisheries Oceanography 28, 532–566.

Taylor, D., Fehon, M., Cribari, K., Scro, A., 2022. Blue crab *Callinectes sapidus* dietary habits and predation on juvenile winter flounder *Pseudopleuronectes americanus* in southern New England tidal rivers. Marine Ecology Progress Series 681, 145–167.

Townsend, D.W., Pettigrew, N., Thomas, M.A., Moore, S., 2023. Warming waters of the Gulf of Maine: The role of shelf, slope, and Gulf Stream water masses. Progress in Oceanography 215, 103030.

Venables, W., Ripley, B., 2002. Modern Applied Statistics with S. Springer.

Wagstaff, M., 2015. Critical forces that structure subtidal eccologial communities in the Gulf of Maine, and the integration of invasive species into these communities, Environmental Biology. University of Massachusetts Boston, Massachusetts, USA, p. 203.

Walter, W., 2021. Green economics: Assessing the feasibility of a New England green crab (Carcinus maenas) fishery through fisherman perspectives. University of New England, Maine, USA, p. 58.

Webber, M., 2013. Results of the one-day green crab trapping survey conducted along the Maine coast from August 27 to 28, 2013, in: Maine Department of Marine Resources (Ed.), Augusta, Maine.

Wells, C., Van Volkom, K., Edquist, S., Marovelli, S., Marovelli, J., 2023. Investigating the impact of introduced crabs on the distribution and morphology of littorinid snails: Implications for the survival of the snail *Littorina saxatilis*. Journal of Experimental Marine Biology and Ecology 569, 151958.

Young, A.M., Elliott, J.A., 2019. Life history and population dynamics of green crabs (*Carcinus maenas*). Fishes 5.

