## Supplemental Materials for "Blue beats green: Agonistic interactions between Atlantic blue crabs and European green crabs in the Gulf of Maine"

**SUPPLEMENTARY MATERIAL**

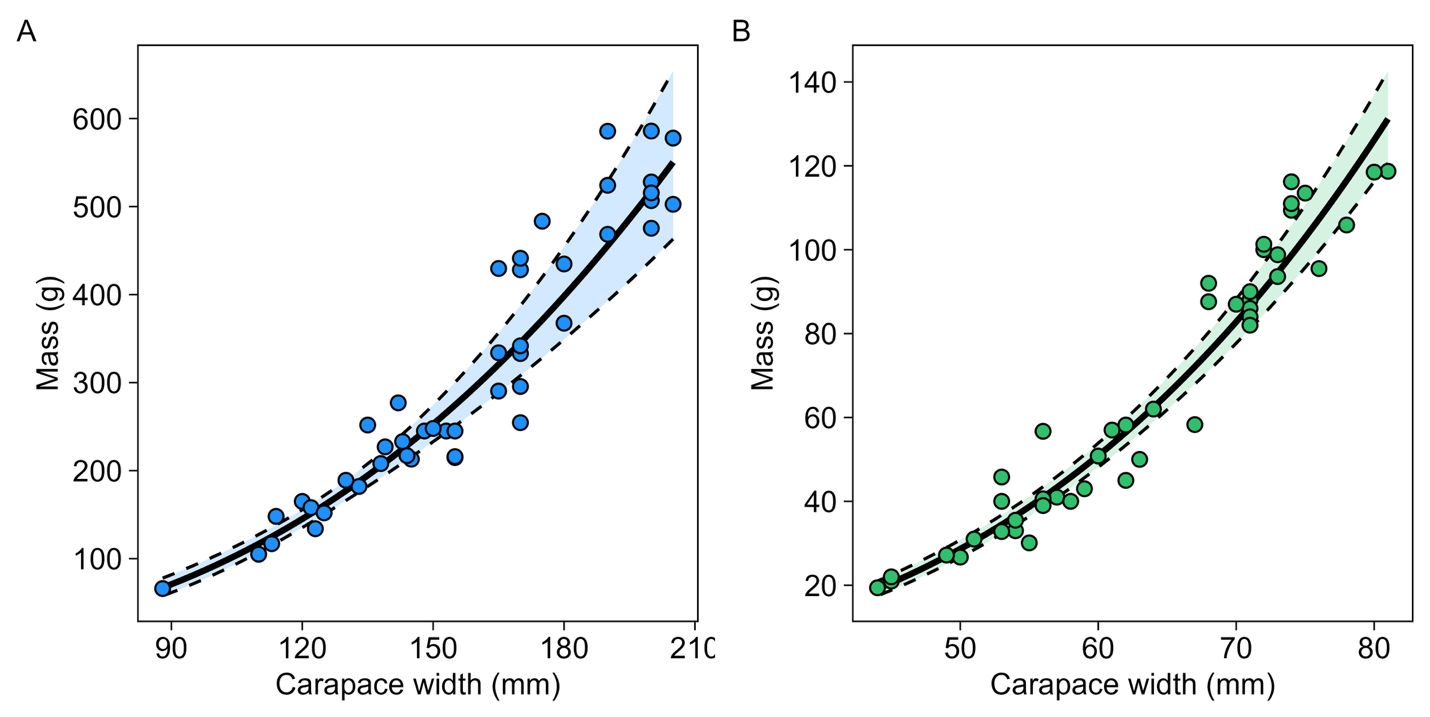

Fig. S1. Relationship between CW and body mass of blue crabs (A) and green crabs (B). For both species, Anova type II tests resulted in statistical significance (*p* < 0.001 for both). Best fit lines are as follows; blue crabs: mass = 7.69 × 10^-4^ × width^2.54^, R^2^ = 0.93; green crabs: mass = 1.25 × 10^-4^ × width^3.16^, R^2^ = 0.95. Bold lines represent model predictions, while shaded regions bound by dashed lines indicate 95% confidence intervals. Points denote observed data.

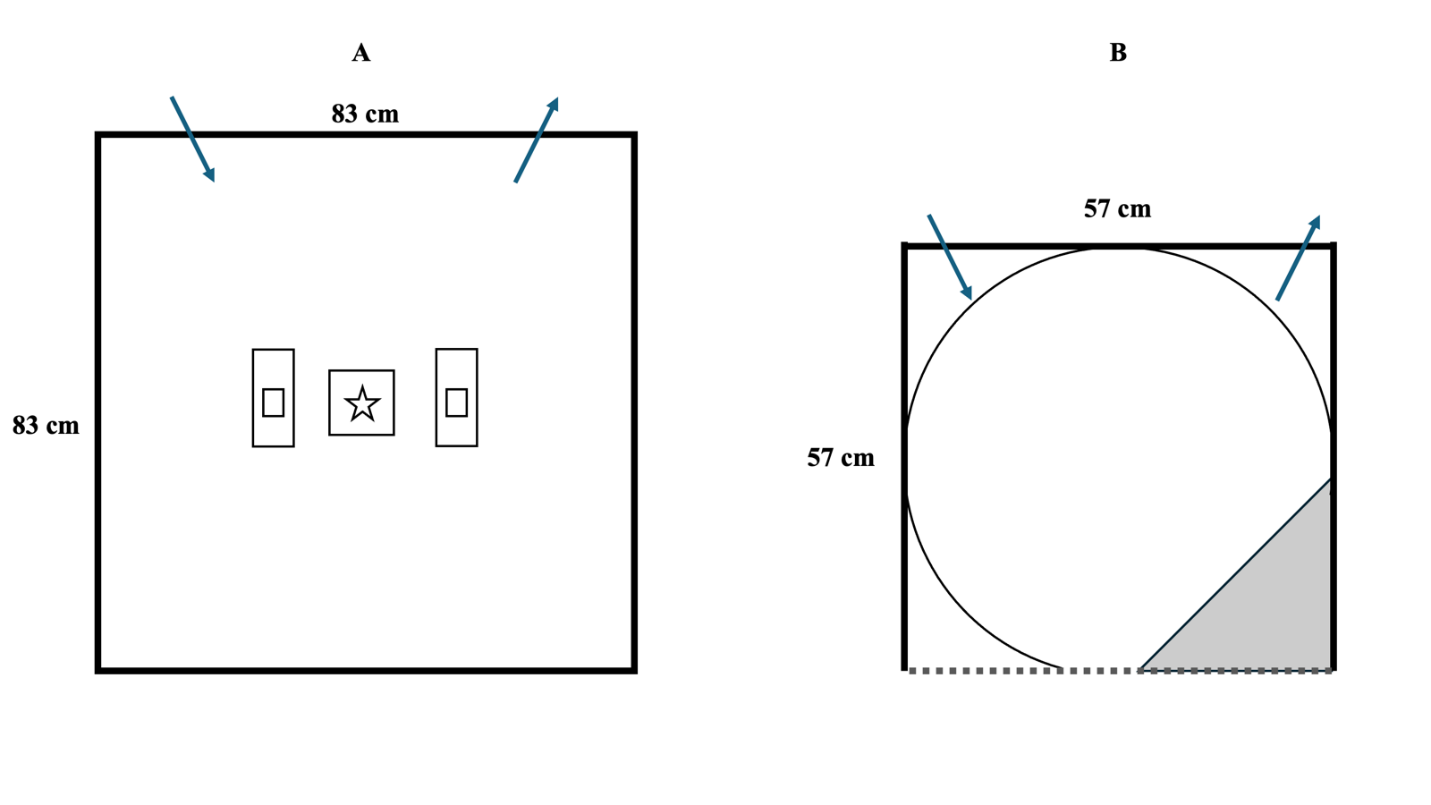

Fig. S2. Tank set up (top-down view) at WNERR (A) and SCSC (B). A rectangle represents a brick, and a star represents a ceramic tile. Water flow is indicated by arrows. Dotted line (B) represents the transparent side through which video was recorded. Shaded region (B) represents the area designated as the corner region.
